# microRNA-210 enhances cell survival and paracrine potential for cardiac cell therapy while targeting mitophagy

**DOI:** 10.1101/2025.01.09.632206

**Authors:** Rita Alonaizan, Ujang Purnama, Sophia Malandraki-Miller, Mala Gunadasa-Rohling, Andrew Lewis, Nicola Smart, Carolyn Carr

## Abstract

The therapeutic potential of presumed cardiac progenitor cells (CPCs) in heart regeneration has garnered significant interest, yet clinical trials have revealed limited efficacy due to challenges in cell survival, retention, and expansion. Priming CPCs to survive the hostile hypoxic environment may be key to enhancing their regenerative capacity. We demonstrate that microRNA-210 (miR-210), known for its role in hypoxic adaptation, significantly improves CPC survival by inhibiting apoptosis through the downregulation of *Casp8ap2*, reduction of caspase activity, and decreased DNA fragmentation. Contrary to the expected induction of Bnip3-dependent mitophagy by hypoxia, miR-210 did not upregulate *Bnip3*, indicating a distinct anti-apoptotic mechanism. Instead, miR-210 reduced markers of mitophagy and increased mitochondrial biogenesis and oxidative metabolism, suggesting a role in metabolic reprogramming. Furthermore, miR-210 enhanced the secretion of paracrine growth factors from CPCs, which promoted *in vitro* endothelial cell proliferation and cardiomyocyte survival. These findings elucidate the multifaceted role of miR-210 in CPC biology and its potential to enhance cell-based therapies for myocardial repair by promoting cell survival, metabolic adaptation, and paracrine signalling.

## Introduction

Clinical efforts to achieve heart regeneration have primarily concentrated on cell therapy, using mesenchymal stem cells, cardiomyocytes or cardiac-committed cells derived from pluripotent stem cells, and presumed cardiac progenitor cells (CPCs). Despite early promise from pre-clinical studies, the administration of these cell populations in clinical trials has shown limited beneficial effects. CPCs have demonstrated the ability to improve cardiac function in animal models of myocardial infarction (MI), despite lacking long-term retention and the ability to generate new cardiomyocytes. It is, therefore, believed that CPCs exert their therapeutic effects primarily in a paracrine fashion, releasing a multitude of bioactive molecules that influence the surrounding microenvironment and promote tissue repair^1^.

CPCs can be isolated and expanded in culture via the formation of cardiospheres to give cardiosphere-derived cells (CDCs), by selecting specific markers such as Kit or Sca1^1^, or by expanding rare clonogenic cells^2^ but this requires an extended period to grow in culture, limiting their immediate clinical applicability as an autologous cell type. The clinical trials using intracoronary administration of autologous CDCs following MI showed a reduction in scar size^3^. Subsequent trials using allogeneic CDCs showed no change in scar size but improved segmental myocardial function post MI^4^ and improved left ventricular ejection fraction in patients with late-stage Duchenne muscular dystrophy^5^. Notable challenges associated with the use of CPCs are the long expansion time, cell death following transplantation and low cell retention. Therefore, it is imperative to establish robust protocols that enable time-effective expansion of CPCs and enhance their survival under ischaemia to fully harness their therapeutic potential in regenerative medicine.

We developed a collagenase-trypsin protocol adapted from studies by Gharaibeh *et al*.^6^ and Okada *et al*.^7^ for isolating slowly adhering cells (SACs) from skeletal muscle. These SACs demonstrated superior differentiation, survival, and therapeutic potential compared to rapidly adhering cells (RACs). When applied to cardiac tissue, the protocol produced a CPC population with more consistent cell yield and expansion time compared to CDCs, while maintaining a similar gene expression profile^8^. We have shown that these CPCs can be induced to differentiate into the cardiomyocyte lineage *in vitro*^9^ using a TGFβ1-based protocol^10,11^. Additionally, we enhanced this differentiation efficiency by stimulating the PPARα pathway through fatty acid supplementation^9^.

One critical factor that plays a pivotal role in enhancing the survival and function of transplanted cells is the cellular response to hypoxia. MicroRNA-210 (miR-210) is a key player in this process. Unique among other hypoxamiRs, miR-210 is consistently and robustly induced under hypoxic conditions, such as those encountered in ischaemic damge^12^. *Casp8ap2,* which encodes caspase 8 associated protein 2, has been shown to be a direct target of miR-210, and has been implicated in cell cycle regulation^13^, transcriptional control^14^ and the activation of caspase 8 during apoptosis^15^. HIF1α has been shown to induce BNIP3-dependent mitophagy as an adaptive metabolic response to hypoxia to increase cell survival^16^, but it is not known if miR-210 acts via this pathway. Upregulating miR-210 in cells before transplantation could significantly enhance their therapeutic efficacy. Therefore, understanding the mechanisms underlying miR-210 activation and exploiting them for therapeutic purposes holds immense promise. The primary aim of this research is to address these critical challenges and pave the way for more effective cell-based therapies.

## Results

### 1. miR-210 improves the survival of CPCs by targeting apoptotic cell death following serum starvation

CPCs were isolated from mouse atria and expanded to passage 7 then transfected with miR-210 or a negative-miRNA. miR-210 overexpression was confirmed 48 and 72 hours after transfection using qRT-PCR (**Figure S1**). To validate miR-210’s pro-survival effect and confirm that it targets cell death pathways in CPCs, expression of its known direct target, *Casp8ap2*^17^, was analysed after miRNA transfection and 72 hours of serum starvation. *Casp8ap2* expression was significantly reduced following miR-210 transfection in comparison to the negative miRNA (**Figure 1A**). A poly-caspase activity assay revealed significantly reduced overall activity of caspases following miR-210 transfection in comparison to the negative miRNA (**Figure 1B**). Finally, reduced apoptotic cell death was confirmed by measuring DNA fragmentation using a TUNEL assay (**Figure 1C-D**).

**Figure 1.**
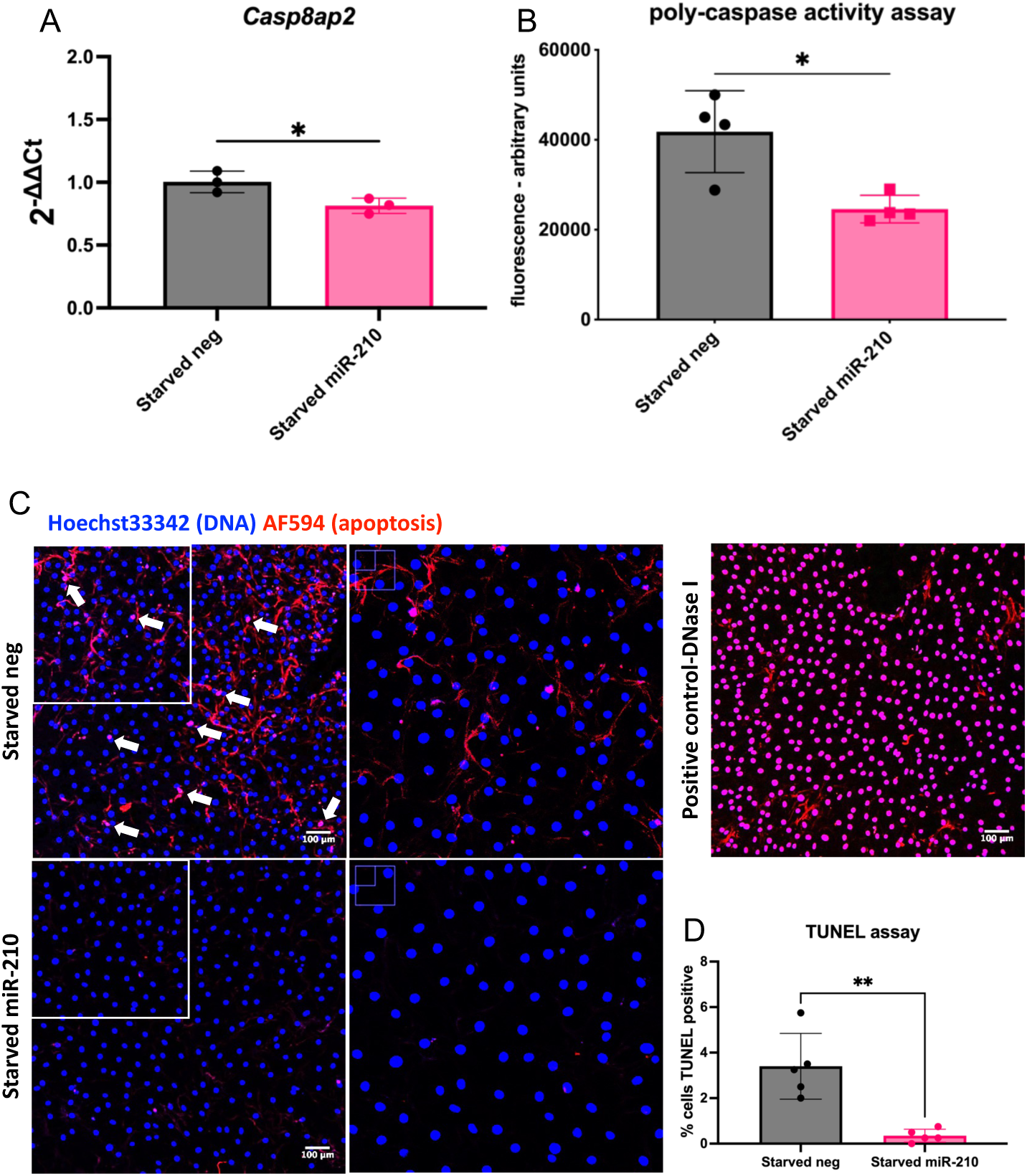
miR-210-transfected CPCs show decreased apoptosis following serum starvation. (A) *Casp8ap2* mRNA levels (n=3) and (B) poly-caspase activity in miR-210-transfected CPCs following serum starvation, in comparison to CPCs transfected with a negative miRNA (n=4). (C) DNA fragmentation represented as TUNEL^+^ cells in miR-210-transfected CPCs following serum starvation, in comparison to CPCs transfected with a negative miRNA. Cells were imaged using an FV1000 confocal microscope. High power inserts are shown by white boxes. DNase treatment was used as a TUNEL assay positive control; scale bar = 100µm. (D) Quantification of the TUNEL assay (n=5). Analysed using a t-test; * p < 0.05, ** p < 0.01.

### 2. Unlike hypoxia, miR-210 does not induce BNIP3 as an anti-apoptotic mechanism

To examine a potential relationship between the hypoxamiR miR-210 and mitophagy-induced cytoprotection, we examined the expression of mitophagy-associated genes in response to hypoxic culture and/or miR-210 overexpression. We found that hypoxic culture increased the expression of *Bnip3*, *Nix* and *Atg4c*, but not *Atg7*, indicating enhanced mitophagy (**Figure S2**).

To assess whether the cytoprotective role of miR-210 acted partly through BNIP3-dependent mitophagy, we knocked down the mRNA levels of *Bnip3* in CPCs using siRNA (**Figure S3**), and measured apoptosis following 72 hours of serum starvation. *Bnip3* knockdown did not decrease miR-210’s capacity to target apoptosis (**Figure 2A-B**). This suggests that unlike the previously described role of HIF1α^16^ miR-210 does not induce BNIP3 as an anti-apoptotic mechanism. A significant reduction in apoptosis was still observed in miR-210-transfected cells in comparison to their respective negative miRNA group in both *Bnip3* siRNA and negative siRNA conditions, which is consistent with miR-210’s pro-survival effect.

**Figure 2.**
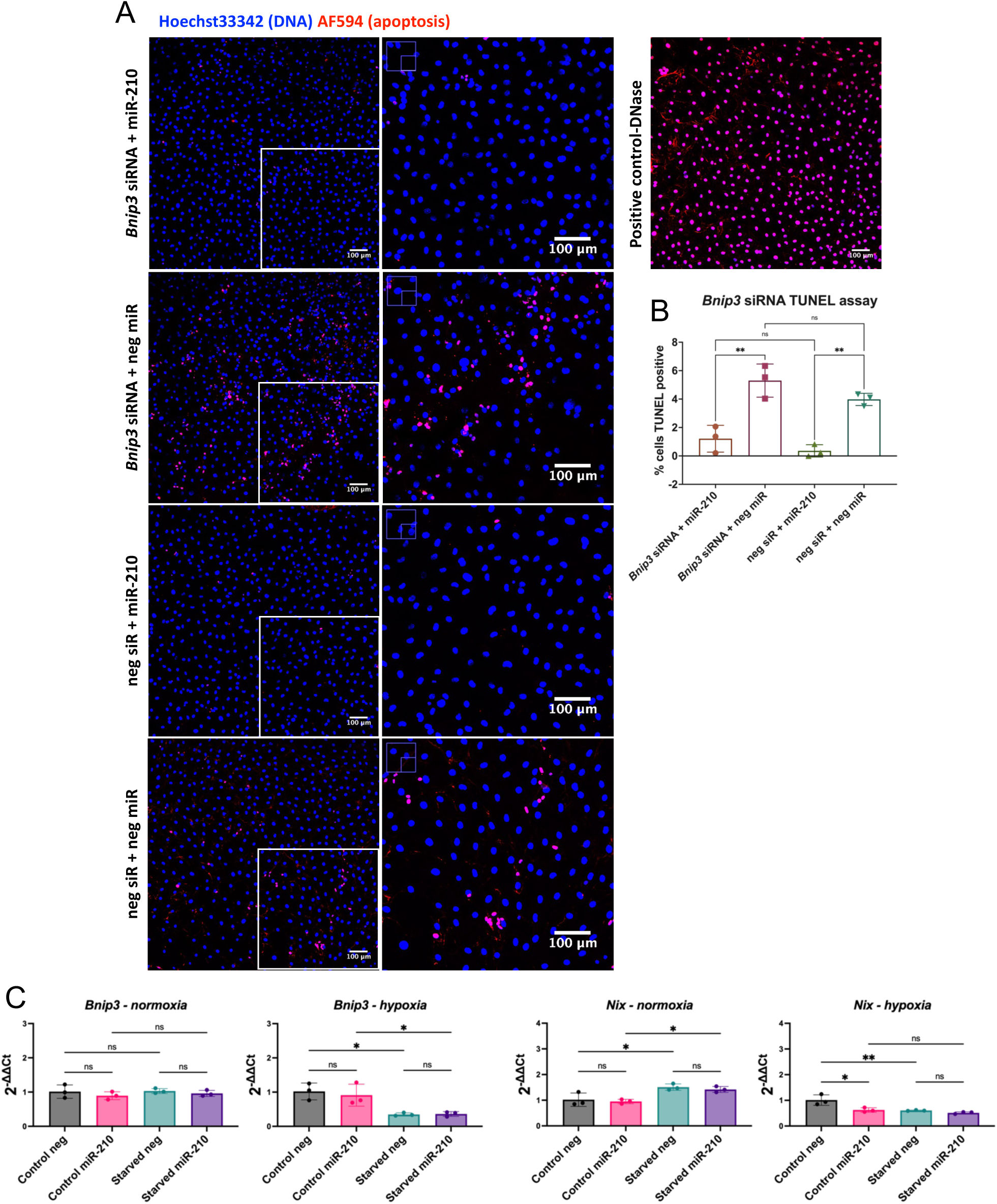
*Bnip3* knockdown in miR-210-transfected CPCs TUNEL assay. (A) DNA fragmentation represented as TUNEL^+^ cells following serum starvation of CPCs transfected with the following: *Bnip3* siRNA+miR-210, *Bnip3* siRNA+negative miRNA, negative siRNA+miR-210, negative siRNA+negative miRNA. Cells were imaged using an FV1000 confocal microscope. High power inserts are shown by white boxes. DNase treatment was used as a TUNEL assay positive control. (B) Quantification of the TUNEL assay. (C) mRNA levels of *Bnip3* and *Nix* in miR-210-transfected CPCs, in comparison to CPCs transfected with a negative miRNA, under the following conditions: control media and normoxia, serum starvation and normoxia, control media and hypoxia, serum starvation and hypoxia. Analysed using a one-way ANOVA with Tukey’s multiple comparisons test (n=3); * p < 0.05, ** p < 0.01.

*Bnip3* and *Nix* mRNA expression was measured in miR-210-overexpressing CPCs cultured under normoxia or hypoxia (**Figure 2C**). We found that miR-210 did not significantly alter *Bnip3* expression, but reduced *Nix* expression in CPCs cultured in control conditions under hypoxia in comparison to their respective negative miRNA group. Interestingly, *Nix* and *Bnip3* expression was decreased in response to serum starvation under hypoxia. In contrast, an increase in *Nix* expression was observed in response to serum starvation under normoxia and no significant change was observed with *Bnip3*. Overall, this shows that, unlike hypoxia, miR-210 does not induce Bnip3-dependent mitophagy as an anti-apoptotic mechanism. In contrast, the reduction in *Nix* levels following miR-210 transfection of CPCs under hypoxia suggested that there might, in fact, be a reduction in mitophagy levels. Therefore, this was further investigated.

### 3. Mitophagy as a target of the hypoxamiR miR-210

Expression levels of *Pink1* were assessed as the PINK1-Parkin pathway is the most widely studied mode of mitophagy. We found that *Pink1* was significantly reduced in miR-210-overexpressing CPCs in comparison to their respective negative miRNA group under all conditions, except for the starved normoxic condition (**Figure 3A**). *Atg4c* expression was reduced in miR-210-overexpressing CPCs in comparison to their respective negative miRNA group in hypoxic conditions whereas *Atg7* expression levels were not significantly altered. We also saw a decrease in the expression of the mitochondrial fission factor *Drp1* in miR-210 overexpressing CPCs in starved cells under normoxia and control cells under hypoxia. A significant increase in *Drp1* levels in response to serum starvation was observed in control and miR-210 over-expressing cells cultured under hypoxia.

**Figure 3.**
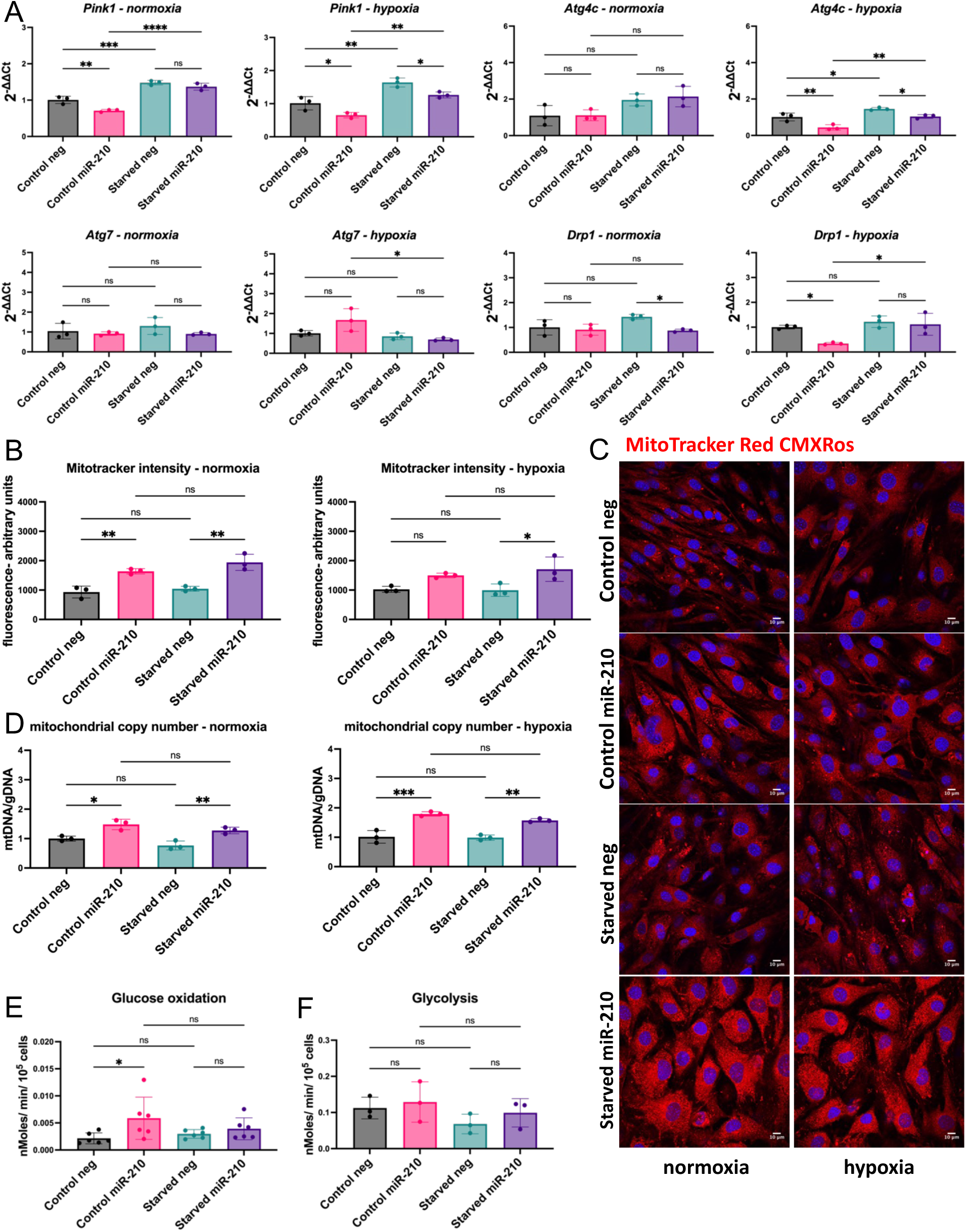
miR-210-transfected CPCs show decreased mitophagy and higher glucose oxidation levels. (A) mRNA levels of *Bnip3*, *Nix*, *Pink1*, *Atg4c*, *Atg7, and* the fission factor *Drp1* in miR-210-transfected CPCs in comparison to CPCs transfected with a negative miRNA, under the following conditions: control media and normoxia, serum starvation and normoxia, control media and hypoxia, serum starvation and hypoxia (n=3); (B-C) Mitotracker intensity quantification and representative images (n=3; scale bar = 10µm); (D) mitochondrial copy number (n=3); (E) glucose oxidation (n=6) and (F) glycolytic (n=3) rates measured as nMoles/ min/ 10^5^ cells, under the following conditions: control media and serum starvation. Analysed using a one-way ANOVA with Tukey’s multiple comparisons test; * p < 0.05, ** p < 0.01, *** p < 0.001, **** p < 0.0001.

Mitotracker CMXRos was used as a measure of mitochondrial membrane potential. We found an increase in Mitotracker CMXRos retention in miR-210-transfected CPCs in comparison to the negative miRNA groups in all groups except control cells under hypoxia. In contrast, there was no significant change in response to serum starvation (**Figure 3B-C**). Mitochondrial copy number was also assessed by mitochondrial DNA/genomic DNA ratio using qRT-PCR. We observed increased mitochondrial copy number in miR-210-transfected CPCs in all conditions in comparison to their respective negative miRNA group (**Figure 3D**).

Mitophagy can act as a modulator of metabolism and cell fate^18^. Additionally, the degree of mitophagy has been shown to be dependent on the metabolic context^19^. We therefore examined the rates of glycolysis and glucose oxidation of miR-210-transfected CPCs to determine if they were consistent with reduced mitophagy. As both measurements involved the use of radioactive material, they were performed in a radioactivity lab where only a normoxic incubator was available for use. We found a significant increase in glucose oxidation rates in control miR-210-transfected CPCs, in comparison to their respective negative miRNA group, but no change after starvation (**Figure 3E**). No significant changes in glycolysis were detected. Altogether, these results suggested reduced mitophagy in response to miR-210 overexpression.

### 4. miR-210 overexpression in CPCs reveals a complex relationship with hypoxia-inducible genes

miR-210 is regarded as an hypoxamiR that is capable of upregulating HIF1α via a positive feedback loop^20^. However, our results indicated that unlike HIF1α, miR-210 may have a role in reducing mitophagy and enhancing oxidative metabolism. This warranted us to further explore the relationship between miR-210 and HIF1α by assessing the effect of miR-210 on the expression of the known HIF1α target pyruvate dehydrogenase kinase 1 (*Pdk1*)^21,22^ as well as targets by which miR-210 has been shown to induce hypoxic adaptations. For example, it has been shown that miR-210 targets *Iscu*, which encodes the mitochondrial iron sulphur scaffold protein, to induce a glycolytic shift in human cancer cell lines^23^. Moreover, we examined the mRNA levels of HIF1α stability regulators *Phd1* and *Phd3*.

We found that *Iscu1* expression was significantly reduced in miR-210-overexpressing CPCs in comparison to their respective negative miRNA group under all conditions. *Iscu2* expression was reduced in miR-210-overexpressing CPCs in comparison to their respective negative miRNA group in starved hypoxic conditions. Moreover, we observed an increase in *Iscu1* and *Iscu2* expression in almost all conditions in response to serum starvation (**Figure 4**). Similar to miR-210 overexpression, hypoxic culture significantly decreased *Iscu1* and *Iscu2* levels (**Figure S4**).

**Figure 4.**
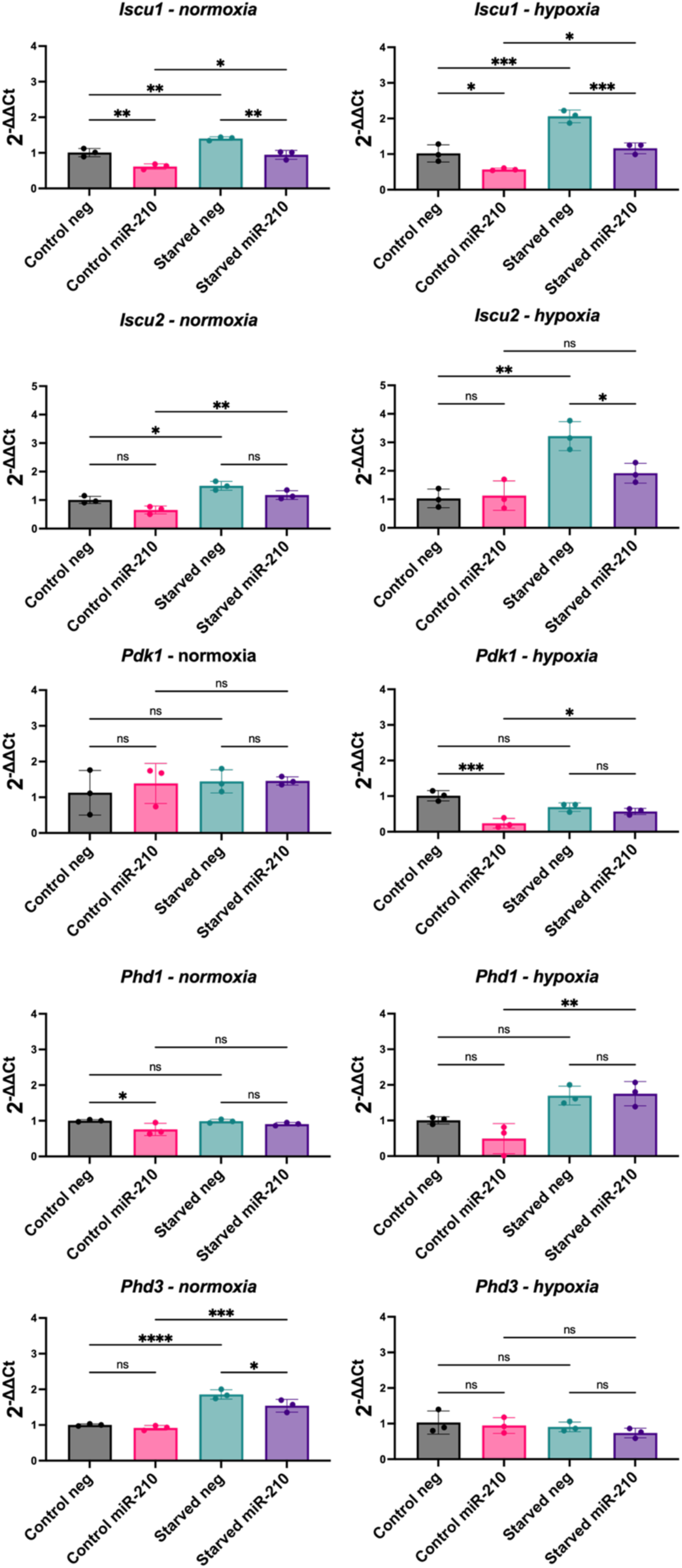
Expression of hypoxia-associated genes in miR-210-transfected CPCs. mRNA levels of *Iscu1*, *Iscu2*, *Pdk1*, *Phd1*, *Phd3* in miR-210-transfected CPCs, in comparison to CPCs transfected with a negative miRNA, under the following conditions: control media and normoxia, serum starvation and normoxia, control media and hypoxia, serum starvation and hypoxia. Analysed using a one-way ANOVA with Tukey’s multiple comparisons test (n=3; * p < 0.05, ** p < 0.01, *** p < 0.001, **** p < 0.0001).

The effect of miR-210 overexpression on *Pdk1*, *Phd1* and *Phd3* was less consistent. *Pdk1* expression was reduced in miR-210-overexpressing CPCs in comparison to their respective negative miRNA group in control media under hypoxia but not in other conditions; *Phd1* was only reduced by miR-210 overexpression in CPCs in control media under normoxia and *Phd3* was only reduced in starved media under normoxia. After serum starvation, we observed increased expression of *Pdk1* and *Phd1* in miR-210-overexpressing CPCs under hypoxia and of *Phd3* expression in both control and miR-210 overexpressing CPCs under normoxia (**Figure 4C**). Unlike miR-210 overexpression, hypoxic culture significantly increased *Pdk1* and *Phd3* expression (**Figure S4**). This revealed a complex relationship between the effect of hypoxamiR miR-210 and that of hypoxia.

### 5. miR-210-overexpressing CPCs enhance HUVEC proliferation and HL-1 cardiomyocyte survival through paracrine mechanisms

Due to the significant implication of paracrine signalling in the therapeutic effect of cell therapy on heart remodelling and regeneration, we assessed the ability of miR-210 to increase the secretion of beneficial growth factors from CPCs. When CPCs were transfected with miR-210, an increase in SCF, IGF-1, and IGFBP-2 secretion was observed compared to the negative groups under normoxia but not hypoxia (**Figure 5A-B**).

**Figure 5.**
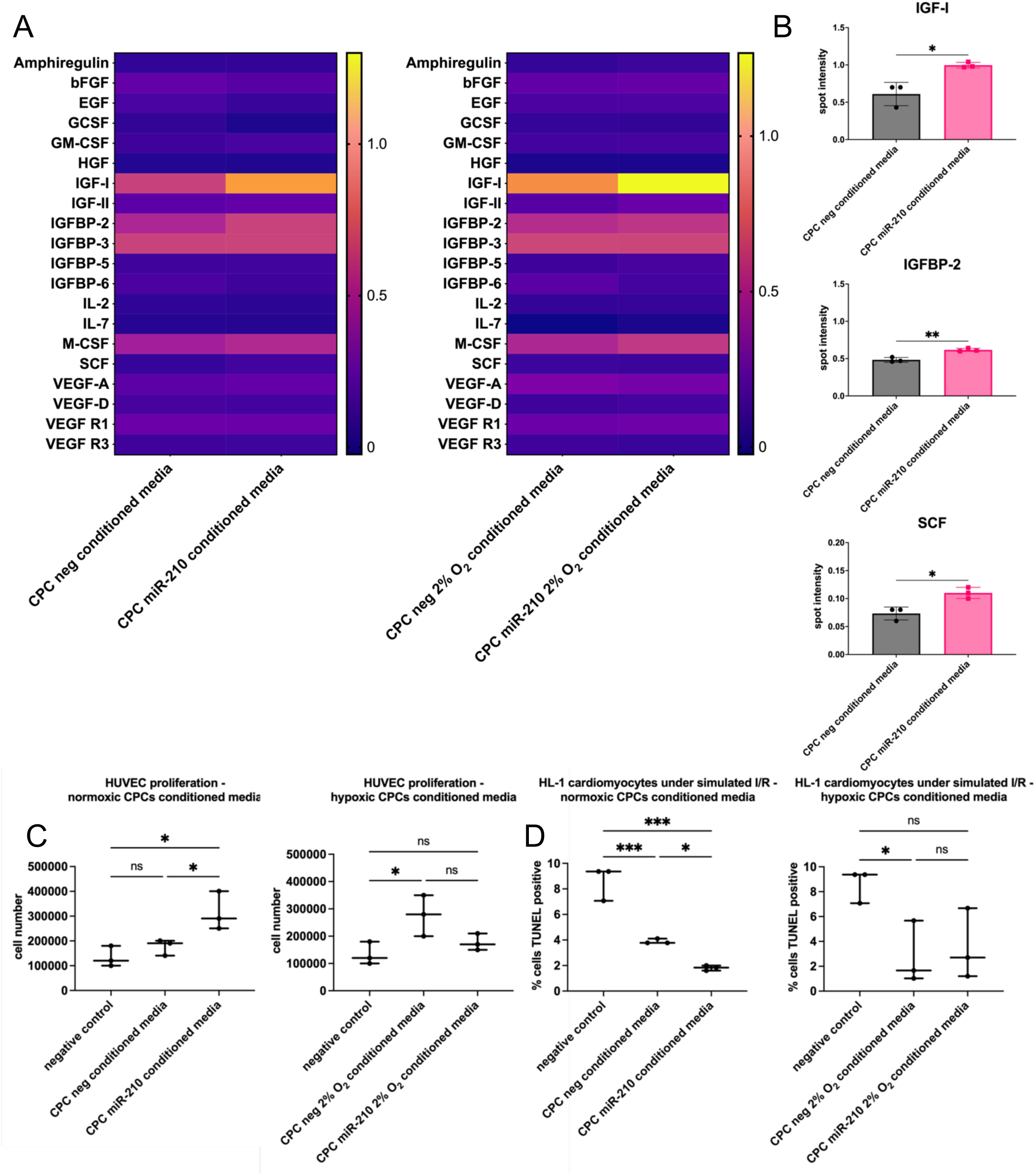
miR-210 enhances the paracrine potential of CPCs as shown by increased HUVEC proliferation and an anti-apoptotic effect in HL-1 cardiomyocytes during indirect co-culture. (A) The secretion of 20 growth factors was assessed in the conditioned media of miR-210-transfected CPCs, in comparison to CPCs transfected with a negative miRNA using an array. Spot intensity of the samples was normalised to the average of the positive control spots of each array. The heatmap illustrates spot intensity (black is low or absent; yellow is high) of the assessed factors secreted from CPCs cultured in normoxia or hypoxia. (B) Individual representation of growth factors that were significantly differentially secreted from CPCs cultured under normoxia. Analysed using a t-test (n=3). (C) HUVEC proliferation after 24-hour culture in 1:1 serum-free HUVEC media and conditioned media or unconditioned media as a negative control. The conditioned media of miR-210-transfected CPCs was compared to that of CPCs transfected with a negative miRNA cultured in normoxia or hypoxia. Analysed using a two-way ANOVA with Tukey’s multiple comparisons test (n=3). (D) TUNEL^+^ cells in HL-1 cardiomyocytes after simulated ischaemia/reperfusion and treatment with CPC conditioned media or unconditioned media as a negative control. The conditioned media of miR-210-transfected CPCs was compared to that of CPCs transfected with a negative miRNA cultured in normoxia or hypoxia. Analysed using a two-way ANOVA with Tukey’s multiple comparisons test (n=3; * p < 0.05, ** p < 0.01, *** p < 0.001).

HUVECs were used to evaluate the paracrine potential of CPC conditioned media and whether this was enhanced by miR-210 transfection. HUVECs were switched to culture in 1:1 serum-free HUVEC media and conditioned media or unconditioned media as a negative control for 24 hours then harvested for cell counting. Conditioned media of CPCs cultured under hypoxia, but not normoxia, enhanced the proliferation of HUVECs in comparison to the negative control. Moreover, conditioned media of normoxic, but not hypoxic, miR-210-overexpressing CPCs significantly enhanced the proliferation of HUVECs in comparison to that of CPCs transfected with the negative miRNA (**Figure 5C**).

We next examined miR-210’s ability to enhance the pro-survival effect of CPC conditioned media on cultured cardiomyocytes. HL-1 cardiomyocytes were subjected to simulated ischaemia/reperfusion (I/R). This was performed by culturing the cardiomyocytes in serum- and glucose-free media and hypoxia (1% O_2_) for six hours to simulate ischaemia, followed by culture in conditioned media or, unconditioned media as a negative control, and normoxia for 24 hours to simulate reperfusion. Conditioned media of CPCs cultured under both normoxia and hypoxia decreased cardiomyocyte apoptosis under simulated I/R in comparison to the negative control (**Figure 5D**). Moreover, conditioned media of normoxic miR-210-overexpressing CPCs further decreased cardiomyocyte apoptosis in comparison to that of CPCs transfected with the negative miRNA.

## Discussion

### miR-210 inhibits apoptotic cell death

Contradictory roles of miRNAs have been described in the literature^24–27^. This may be attributed to the cell type and its gene expression profile and cellular microenvironment. It may also be a result of regulatory loops by the miRNA’s own targets as a miRNA and its targets can form both feedback and feedforward loops^28,29^. Moreover, the ability to have multiple targets in various pathways is a miRNA characteristic, which may give them the potential to conduct different regulatory effects.

In this study, we showed that miR-210 has an anti-apoptotic effect in CPCs *in vitro* by assessing *Casp8ap2* expression, caspase activation and DNA fragmentation. We found that serum-starved CPCs showed significantly decreased *Casp8ap2* mRNA levels, poly-caspase activity and DNA fragmentation when transfected with miR-210 in comparison to the negative miRNA indicating reduced apoptosis and superior survival potential. Although a pro-apoptotic role for miR-210 via ROS induction has been described^30^, miR-210 has been shown to inhibit apoptotic cell death by targeting *Casp8ap2*^17^, *Ptp1b* and *Dapk1*^31^. Moreover, miR-210 promoted cardiomyocyte proliferation and inhibited apoptosis by targeting *Apc*, which encodes a cell cycle inhibitor^32^. miR-210 has also been shown to be enriched in exosomes secreted by CDCs and these exosomes have been shown to have a cytoprotective role^33^. Moreover, miR-210 is upregulated in KIT^+^ cell-derived exosomes subjected to hypoxia. These hypoxic exosomes improved cardiac function in a rat model of cardiac injury^34^.

### miR-210 has a role in the regulation of mitochondrial fission and autophagy

A role for miR-210 in mitophagy has not been previously explored. However, our results showing decreased *Nix*, *Pink1*, and *Drp1* levels, and upregulated mitochondrial copy number and MitoTracker Red CMXRos retention in miR-210-transfected CPCs support a role for miR-210 in mitophagy. A study in endothelial progenitor cells revealed that transfecting the progenitors with miR-210 enriched for miR-210 in their secreted exosomes^35^. Treatment with these exosomes of endothelial cells subjected to I/R injury increased ATP levels and mitochondrial membrane potential and decreased mitochondrial fission^35^. This raises the question of whether mitophagy is a direct or indirect target of miR-210. A study reported that fibroblasts lacking caspases 3 and 7 are resistant to loss of mitochondrial membrane potential^36^. Therefore, it can be argued that miR-210-induced caspase inhibition may be upstream of mitochondrial events. However, in our study, the reduction of mitophagy in miR-210-transfected CPCs was not confined to conditions of serum starvation where caspase activation was higher than control conditions.

Mitophagy has been classically viewed as a mitochondrial quality control mechanism but multiple reasons for mitochondrial elimination have been described. This is supported by observations ranging from programmed forms of mitophagy to basal mitophagy in different tissues demonstrating the complex and multi-factorial nature of mitophagy^37^. The levels of basal activation are heterogenous across tissues, and across different cells within the same tissue, highlighting context-dependant regulation that is not yet understood^19^. Indeed, it has been shown that levels of basal mitophagy depend on the metabolic context of the cell where greater levels are observed in highly metabolic cells such as cardiomyocytes and dopaminergic neurons^19^. Therefore, the observed downregulation in mitophagy is not necessarily linked to the pro-survival role of miR-210 in CPCs.

Moreover, programmed mitophagy describes instances where mitochondria are eliminated for developmental, physiological or metabolic purposes^38^. Cell fate determination and differentiation often involve metabolic reprogramming that implicate mitophagy. A requirement for mitophagy to eliminate mitochondria is observed during the reprogramming of somatic cells into pluripotent stem cells which are highly glycolytic^39–41^. In this study, we observed an increase in glucose oxidation in miR-210-transfected CPCs cultured in control conditions under normoxia concordant with reduced mitophagy. These results, in addition to the previous data, further support the notion that the observed downregulation in mitophagy is not necessarily linked to the pro-survival role of miR-210 and may be a distinct function to induce a metabolic switch.

It remains ambiguous why an hypoxia-inducible miRNA would upregulate mitochondrial networks when the lack of oxygen would stimulate a HIF1α-induced metabolic switch to upregulate glycolysis and mitophagy^16,37^. Studies have reported that miR-210 has a negative effect on mitochondrial respiration with direct targets including *Iscu1/2* and *Cox10*, which are important factors in the mitochondrial electron transport chain^42–45^. Indeed, we found a downregulation in *Iscu1/2* mRNA levels in miR-210-transfected CPCs, similar to the downregulation observed in CPCs cultured under hypoxia. However, when we examined *Pdk1* and *Phd1/3,* we found opposing effects of miR-210 transfection and hypoxic culture on their mRNA levels. PHDs are oxygen sensors that use oxygen as a co-substrate to induce the hydroxylation of HIF1α and subsequent polyubiquitination and proteasomal degradation^46^. Hypoxia, in turn, induces PHDs to allow for rapid HIF1α degradation upon reoxygenation^47–49^. Moreover, HIF1α upregulates PDK1 to inactivate pyruvate dehydrogenase, thereby blocking the entry of pyruvate into the mitochondria for oxidative phosphorylation^48^. In contrast, we saw a decrease in *Pdk1* and an increase in glucose oxidation, in miR-210-overexpressing CPCs under normoxia. Lethal forms of mitophagy have been described, for example, ceramide mediates caspase-independent cell death via excessive activation of mitophagy^50^. Therefore, miR-210 may be downregulating mitophagy to protect against lethality of excessive mitophagy that may occur in hypoxic environments.

Changes observed as a result of serum starvation include a consistent increase in *Iscu1/2* mRNA levels; and an increase in *Pdk1*, *Phd1* and *Phd3* in some conditions. These observed changes in response to serum starvation are contradictory as both components involved in inducing an hypoxic phenotype (*Pdk1*) and others involved in inducing oxidative metabolism (*Iscu1/2*) are upregulated. Moreover, the upregulation in *Phd1/3* mRNA levels following serum starvation is similar to that observed following hypoxic culture. These changes do not appear to be differentially modulated in different conditions (i.e. neg-vs. miR-210-transfected CPCs or CPCs under normoxia vs. hypoxia). It is unclear why these various components would be upregulated following serum starvation, although this may explain why no significant changes were observed in the metabolic rates of CPCs following serum starvation. It has been previously shown that serum starvation alone induces mitochondrial fission^51^, whereas eliminating glutamine or amino acids induces mitochondrial fusion suggesting that starvation conditions can differentially affect mitochondrial fission and fusion^52^. Moreover, human fibroblasts subjected to serum starvation showed a continuous decrease in cellular ATP levels^53^. Serum starvation has been shown to induce HIF1α as a pro-survival factor, even in normoxic cells^54,55^. All these reports are consistent with enhanced mitophagy and increased *Pdk1* and *Phd1/3* expression, but contradict the observed increase in *Iscu1/2* mRNA levels. The contradictory effects of serum starvation may explain why the benefits of miR-210 transfection were more apparent in control conditions as opposed to starved conditions.

We examined the effect of miR-210 overexpression on *Bnip3* and *Nix* as they are both direct targets of HIF1α^56,57^. BNIP3 has been reported to induce apoptosis, necrosis, or autophagy depending on the cellular context^58^. The cytoprotective role of BNIP3-dependent mitophagy has been demonstrated in human primary epidermal keratinocytes^59^. Interestingly, *Nix* expression was increased in response to serum starvation under normoxia, but both *Nix* and *Bnip3* expression showed an opposite direction of change following serum starvation under hypoxia. BNIP3 and NIX share high sequence similarity and identical functionality has been described. They are therefore expected to follow the same pattern. Both genes are induced by HIF1α^56,57^ but their expression needs to be tightly regulated as over expression induces cell death^60^. Hypoxic culture increased *Bnip3* and *Nix* mRNA levels and showed a cytoprotective role in CPCs. Therefore, the opposite direction of change in *Bnip3* and *Nix* mRNA levels following serum starvation under hypoxia in comparison to normoxia may be a mechanism by which hypoxia fine-tunes their levels to protect against cell death.

Overall, our results demonstrated a pro-survival role for miR-210 in CPCs by showing that it reduced apoptosis following serum starvation. Our results also indicated a role for miR-210 in downregulating mitophagy, which is contrary to hypoxia but may be a mechanism by which miR-210 induces a metabolic shift and/or protects against lethality of excessive mitophagy in hypoxia. Whether or not the induced upregulation in mitochondrial dynamics by miR-210 is beneficial to the survival potential of transplanted CPCs in ischaemic heart tissue may be revealed by *in vivo* cell survival and retention studies. miR-210-transfected MSCs showed enhanced engraftment at four days after transplantation in infarcted rat hearts^61^. Moreover, we found that miR-210 reduced the expression of its known direct targets, *Iscu1/2*, which are also downregulated in hypoxia. This may be an indication of the capability of miRNAs to conduct different regulatory functions due to their ability to have multiple targets in various pathways. Our findings therefore highlight that the relationship between miR-210, HIF1α and hypoxia is not as straightforward as once thought and warrants further investigation.

### miR-210 enhances cell paracrine potential

miR-210 improved CPC paracrine function by increasing SCF, IGF-1 and IGFBP-2 secretion. SCF is a direct target of HIF1 and HIF2 and has been implicated in angiogenesis^62–65^. Cardiomyocyte-specific overexpression of SCF decreased myocardial apoptosis and improved cardiac function following MI^66^. Both IGF-1 and IGFBP-2 have been implicated in promoting angiogenesis. IGF-1 has also been shown to have a cytoprotective effect in the infarcted heart. This is concordant with the angiogenic effect of miR-210. Injection of a miR-210 vector into infarcted mouse hearts increased capillary density^31^ and miR-210 has been demonstrated to target and repress *Efna3*, which encodes an inhibitor of angiogenesis^67^. *Efna3* repression by miR-210 in HUVECs is essential for the initiation of HUVEC tubulogenesis, revealing its vital role in angiogenesis^67^. Moreover, miR-210 overexpression in HUVECs stimulated cell migration and tubulogenesis, possibly through a Notch-dependant mechanism^68^. Our results show that miR-210 may have a role in regulating angiogenesis and/or cell survival by targeting IGF and SCF signalling. We observed an increase in the proliferation of HUVECs following incubation in the conditioned media of CPCs cultured under hypoxia, but not normoxia, compared to the negative control. This is similar to the effect of CDC conditioned media on HUVEC tubulogenesis^69^ and studies reporting increased angiogenic paracrine signalling following hypoxic preconditioning of a variety of cell types^70–73^.

Moreover, conditioned media of normoxic miR-210-overexpressing CPCs enhanced the proliferation of HUVECs in comparison to that of CPCs transfected with the negative miRNA, which may also be explained by an increase in the levels of pro-survival factors in the secretome of miR-210-transfected CPCs. We found that conditioned media of CPCs cultured under both normoxia and hypoxia decreased cardiomyocyte apoptosis under simulated I/R in comparison to the negative control. Likewise, a cytoprotective role of SCA1^+^ cell conditioned media has been demonstrated following hypoxic injury in HL-1 cardiomyocytes^74^. Moreover, we found that the conditioned media of normoxic miR-210-overexpressing CPCs further decreased cardiomyocyte apoptosis in comparison to that of CPCs transfected with the negative miRNA. miR-210 has been shown to transfer to primary neonatal cardiomyocytes co-cultured with MSCs via Cx43 gap junctions, resulting in reduced expression of its pro-apoptotic target *Casp8ap2* and improved survival^61^. Moreover, injection of a miR-210 vector into infarcted mouse hearts reduced the percentage of TUNEL^+^ cardiomyocytes^31^. This observation may therefore be due to the secretion of miR-210 itself or the effect of other proteins or genetic material that may have been upregulated in the CPC secretome.

In conclusion, this study provides insights into the use of miR-210 to enhance the therapeutic potential of cell populations for transplantation following cardiac injury, and the relationship between miR-210 and mitophagy that open the door for further studies to uncover the behaviour of miR-210 as an hypoxia miRNA.

## Methods

### 1. Mice

C57BL/6 mice (Harlan, Oxon, UK) were kept under in a 12-hour light-dark cycle, and controlled conditions of temperature, humidity, with free access to water and chow. All animal procedures were reviewed and approved by the University of Oxford Animal Welfare and Ethical Review Board and conform to the Animals (Scientific Procedures) Act 1986 incorporating Directive 2010/63/EU of the European Parliament.

### 2. Isolation and expansion of mouse CPCs

Mice were terminally anesthetised with isoflurane and hearts were isolated and washed with Dulbecco’s phosphate buffered saline (DPBS) containing 50mg of primocin (antimicrobial agent; InvivoGen). Mouse atrial appendages were dissected and minced mechanically into 1 - 2 mm^3^ pieces. The tissue pieces were then transferred into a digestion mix (0.1% trypsin and 0.1% Collagenase II (Calbiochem, 286U/mg) in DPBS) and incubated in a water bath at 37°C for a total of 40 minutes. Every 10 minutes, the digestion mix was mechanically triturated by pipetting, left on ice for 1 minute to settle and the supernatant was collected. Fresh digestion mix was added again to the tissue pieces for a total of 4 digestions. At the final digestion step, sterile 19G needles and 1ml syringes were used to triturate the tissue. After each digestion, the supernatant was neutralised, and the cell suspension was resuspended in fresh complete explant medium (CEM)^75^ to be plated in a 12-well plate through a 40µm cell strainer. The wells were pre-coated with 50µl per cm^2^ of fibronectin bovine plasma (2µg/ml in DPBS; Sigma-Aldrich) and cells were allowed to attach for 3 days without media changes.

### 3. Cell culture

Primary CPCs were cultured in IMDM (ThermoFisher) supplemented with 20% FBS, 100U/ml penicillin, 100μg/ml streptomycin and 2mM L-glutamine (ThermoFisher) and plated on fibronectin-coated flasks. HL-1 cardiomyocytes^76^ were maintained in Claycomb medium (Sigma-Aldrich), supplemented with 100U/ml penicillin, 100μg/ml streptomycin and 2mM L-glutamine (ThermoFisher), 100µM norepinephrine (Sigma-Aldrich) in 30mM L-ascorbic acid (Sigma-Aldrich), and 10% FBS and plated on flaks pre-coated with 0.02% (wt/vol) gelatin (Sigma-Aldrich) containing 5µg/ml fibronectin. Primary HUVECs, obtained from Lonza (USA), were maintained in complete EGM-Plus (Cell Biologics – USA) supplemented with the provided supplement BulletKit: Endothelial Growth Supplement, L-glutamine, ascorbic acid, hydrocortisone hemisuccinate, human epidermal growth factor, heparin, gentamicin/amphotericin-B, and 2% FBS.

### 4. Indirect co-culture of CPCs with HL-1 cardiomyocytes and HUVECs

CPC conditioned media was collected after 48 hours of culture in serum-free IMDM. HUVECs were seeded at equal densities in 12-well plates and allowed to attach overnight in EGM-Plus media then subjected to serum starvation by switching to culture in 1:1 serum-free EGM-Plus media and CPC conditioned media or unconditioned media as a negative control for 24 hours before harvesting for cell counting. HL-1 cardiomyocytes were seeded at equal densities on gelatin-coated coverslips in 24-well plates and allowed to attach overnight in Claycomb media then subjected to simulated ischaemia/reperfusion (I/R). This was performed by culturing the cardiomyocytes in serum- and glucose-free RPMI and hypoxia (1% O_2_) for six hours to simulate ischaemia. HL-1 cardiomyocytes were then switched to culture in 1:1 serum-free RPMI media and conditioned media or, unconditioned media as a negative control, and normoxia for 24 hours to simulate reperfusion then assessed for apoptotic cell death using a TUNEL assay.

### 5. Immunocytochemistry

Cells were seeded on cover glasses in 24-well plates and allowed to attach prior to fixation with 4% PFA (Santa Cruz) for 30 minutes at 4°C. For intracellular staining, cells were permeabilised with 0.5% Triton X-100 (Sigma-Aldrich) for 10 minutes at RT and washed twice with DPBS. Cells were incubated in blocking solution (2% BSA, 2% FBS ± 0.05% Triton X-100) for 1 hour at RT. Cells were incubated with the primary antibody in blocking solution overnight at 4°C then washed with DPBS. The secondary antibody was added for 30 minutes at RT then slides were washed with DPBS and nuclei were stained with DAPI for 10 minutes at RT. Samples were mounted with 1:1 DPBS:Glycerol and scanned using an Inverted Olympus FV1000 Confocal system. Acquired images were processed using Fiji software.

### 6. RNA extraction and qRT-PCR

RNA was extracted from frozen cell pellets, using the RNeasy Mini Kit (Qiagen) and cDNA was synthesised, using the High-Capacity cDNA Reverse Transcription kit (ThermoFisher) according to manufacturer’s protocol. Real time quantitative PCR was performed using the Applied Biosystems StepOnePlus Real-Time PCR System (Applied Biosystems, ThermoFisher). Relative mRNA levels were normalised to the housekeeping gene *Hprt*. SYBR Green fluorescent intercalating dye was used as a detection system for the reaction (SYBR Green PCR mastermix, ThermoFisher). For miR-210 qRT-PCR, RNA was extracted using miRNeasy Mini Kit (Qiagen) and cDNA was synthesised using miScript II RT Kit (ThermoFisher). Relative miR-210 levels were normalised to the housekeeping gene *Snord68* using the miR-210_2 and SNORD68_11 miScript Primer Assays. miScript SYBR Green PCR Kit (ThermoFisher) was used as a detection system for the reaction. Relative quantification of target gene expression was performed using the ΔΔCt Livak method^77^, plotting the data as 2^-ΔΔCt±SE^. Primer information is provided in supplementary material (S.Table 1).

### 7. Quantifying mitochondrial copy number by qPCR

DNA was extracted from cells using the DNeasy Blood & Tissue Kit (Qiagen) and SYBR Green fluorescent intercalating dye was used as a detection system for the reaction (as described in table 2.2). The gDNA:mtDNA ratio was calculated by normalising the mitochondrial gene NADH dehydrogenase 1 (*mt-Nd1*) to the nuclear gene beta-2 microglobulin (*B2m*) using the ΔΔCt Livak method^77^, plotting the data as 2^-ΔΔCt±SE^.

### 8. miRNA and siRNA transfection using DharmaFECT

Cells were seeded and allowed to attach overnight. DharmaFECT Transfection Reagent #1 (ThermoFisher) was used according to manufacturer’s instructions. Briefly, miRNA or siRNA oligonucleotides and DharmaFECT were diluted in antibiotics/serum-free IMDM separately and incubated for 5 minutes at RT. The two solutions were then combined, mixed by gentle pipetting, and incubated for 20 minutes at RT before adding antibiotics-free CEM to the mix. The final concentration of oligonucleotides was 100nM, unless otherwise indicated. Cells were incubated with the DharmaFECT-oligonucleotide solution for 48 hours. The miRNAs and siRNAs used were: mmu-miR-210-3p mirVana miRNA mimic ID: MC10516 (#4464066), mirVana miRNA Mimic Negative Control (#4464059), Silencer Select Pre-Designed siRNA Application Silencer Select Assay ID: S63060 (#4390771), Silencer Select GAPDH Positive Control siRNA (#4390849), Silencer Select Negative Control No. 2 siRNA (#4390846) (ThermoFisher).

### 9. FAM-FLICA Poly Caspase activity assay

The FAM-FLICA Poly Caspase activity kit measures apoptosis by detecting active caspases in cells via a cell-permeable reagent called the Fluorochrome Inhibitor of Caspases (FLICA), which comprises a caspase inhibitor sequence linked to a green (Carboxyfluorescein, FAM) fluorescent label. Cells were plated in Falcon 96-well black microplates with clear flat bottom and allowed to reach 70-80% confluency. 30X FAM-FLICA reagent was added at 1:30 after experimental manipulation of the plated cells, incubated for 45 minutes at 37°C while mixing gently every 10 minutes, washed with media and incubated for 60 minutes at 37°C allowing unbound excess FLICA to diffuse out of the cells. Cells were washed with fresh media and endpoint fluorescence was measured using an excitation wavelength of 488nm and emission filter of 530nm on the Fluostar OPTIMA Microplate Reader (BMG Labtech, Germany).

### 10. Terminal deoxynucleotidyl transferase dUTP nick end labelling assay (TUNEL) assay

The TUNEL assay was performed using Click-iT TUNEL Alexa Fluor 594 Imaging Assay (ThermoFisher), as per the manufacturer’s instructions. DNase-I treated slides were used for positive control. Samples were imaged using an Inverted Olympus FV1000 Confocal system. The percentage of cell apoptosis was determined by quantifying cells visualised in 3 random fields. Acquired images were processed using the Fiji software plugin JACoP^78^.

### 11. MitoTracker Red CMXRos staining

A working solution of 200nM MitoTracker Red CMXRos, in basal IMDM, was added to the cells and incubated for 40 minutes. Cells were washed with DPBS and fixed with 4% PFA for 10 minutes at RT before staining with DAPI and imaged using an Inverted Olympus FV1000 Confocal system at the excitation/emission wavelengths of 579/599 nm. Fluorescence intensity was quantified using Fiji software by assessing cells visualised in 3 random fields and measuring stained and background areas. Corrected total cell fluorescence (CTCF) was calculated using the following formula^79^: CTCF = integrated density – (area of selected cell × mean fluorescence of background)

### 12. Growth factor array

The conditioned media was assessed by a RayBiotech C-Series Mouse Growth Factor Antibody Array Kit that detects 20 growth factors, many of which had been implicated in cardiac repair. CPCs were transfected with either a negative miRNA or miR-210, then cultured under normoxia or hypoxia on fibronectin-coated plates in basal IMDM (as required by the manufacturer). Culture medium was collected after 48 hours and stored at −80°C. Before use, culture medium was spun down to remove cell debris then run undiluted on the array following manufacturer’s instructions. The arrays were imaged using a LI-COR scanner. The spots were analysed using a pixel counting programme (ImageStudioLite) and the pixel counts normalised to the average of the positive control spots of each array.

### 13. ^14^C Glucose oxidation and ^3^H glycolysis measurements

Glucose oxidation was measured using the method of Collins *et al*.^80^, with some modifications as described in Malandraki-Miller *et al*.^9^, which is dependent on the release of CO_2_ during the passage of acetyl-CoA through the citric acid cycle. CPCs were seeded on fibronectin-coated 24-well plates and the experimental treatment was carried out. Subsequently, CPCs were washed with DPBS then incubated for 3 hours in no-glucose basal IMDM (ThermoFisher) supplemented with radiolabelled glucose at 12mM containing 0.185 MBq D-U-^14^C-glucose (PerkinElmer). Wells with IMDM alone were assessed as negative controls and wells without cells containing IMDM with ^14^C-glucose were used to measure background radioactivity. The ^14^CO_2_ produced by the glucose oxidation was trapped using 40% KOH-soaked filter papers placed in the wells of a 24-well plate used as a lid of the apparatus. A perforated rubber gasket, with holes corresponding to each well, separated the two plates. To ensure the system was sealed and avoid ^14^CO_2_ leakage, the plates were wrapped tightly with Parafilm and screwed into a metal frame. 70% perchloric acid was added to the wells to kill the cells and trigger the release of ^14^CO_2_ after a 2-hour incubation. It was injected through the top holes in the top plate, through the seal and into the well. The holes were instantly resealed with new parafilm. Subsequently, the samples were allowed to sit for 1 hour to allow the complete release of ^14^CO_2_ that was dissolved in the media as HCO^-^_3_. The filter papers containing trapped ^14^CO_2_ were placed in scintillation vials filled with 10mls of scintillation fluid (MP Biomedicals) and analysed using the Tri-Carb 2800TR Liquid Scintillation Analyzer, which measured radiation counts per minute. Glycolysis was measured as described in Malandraki-Miller *et al*.^9^. CPCs were incubated for 6 hours in no-glucose basal IMDM supplemented with radiolabelled glucose at 12mM containing 0.118 MBq ^3^H-glucose (PerkinElmer). Wells with IMDM alone were assessed as a negative control. Following incubation, the culture media contained both released ^3^H_2_O and remaining ^3^H-glucose. Therefore, ^3^H_2_O was separated from ^3^H-glucose using a Dowex 1×4 chloride form (Sigma-Aldrich) anion exchange column, which binds the ^3^H-glucose and elutes ^3^H_2_O. To prepare the columns, 250g of Dowex resin was added to a 1.25M NaOH and 1.61M boric acid solution and washed with distilled H_2_O until the pH reached 7.5. The prepared solution was then added to glass Pasteur pipettes plugged with glass wool. Scintillation vials were filled with 10mls of scintillation fluid (MP Biomedicals) and placed underneath the columns to collect the eluted samples. Subsequently, 200μl of each culture media sample was added to a column allowing for the ^3^H-glucose to bind to the column for 15 minutes before washing each column with 2mls of distilled H_2_O for ^3^H_2_O to be eluted into the scintillation vials.

The samples were analysed using the Tri-Carb 2800TR Liquid Scintillation Analyzer (PerkinElmer), which measured radiation counts per minute.

### 14. Statistical analysis

Results are presented as mean ± standard deviation. All statistical analysis was performed using Graphpad Prism. Data were analysed using unpaired two-tailed Student’s t-test or one-way analysis of variance (ANOVA) unless otherwise indicated. Statistical significance was accepted at p < 0.05.

## Author contributions

Conceptualisation: RA, NS, CC. Investigation: RA, UP, SMM, MGR, AL. Funding acquisition: RA. Writing: RA. Editing: NS, CC.

## Acknowledgements

This work was supported by the King Faisal Specialist Hospital & Research Centre (PhD scholarship to RA).

## Supplementary Material

**Figure S1.**
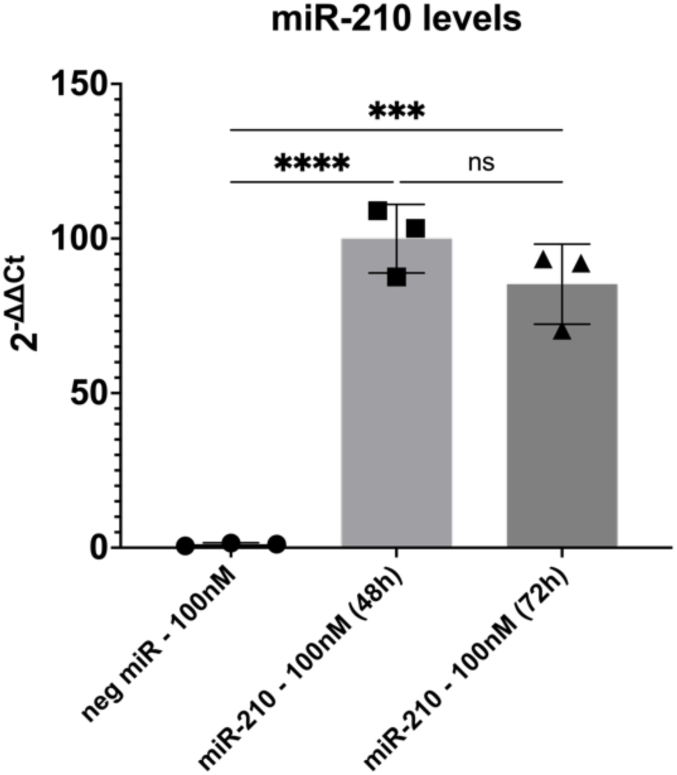
Confirmation of miR-210 overexpression after transfection in CPCs. miR-210 levels in CPCs measured by qRT-PCR at 48 and 72 hours following DharmaFECT transfection with miR-210 compared to CPCs transfected with the negative miRNA. Analysed using a one-way ANOVA with Tukey’s multiple comparisons test (n=3).

**Figure S2.**
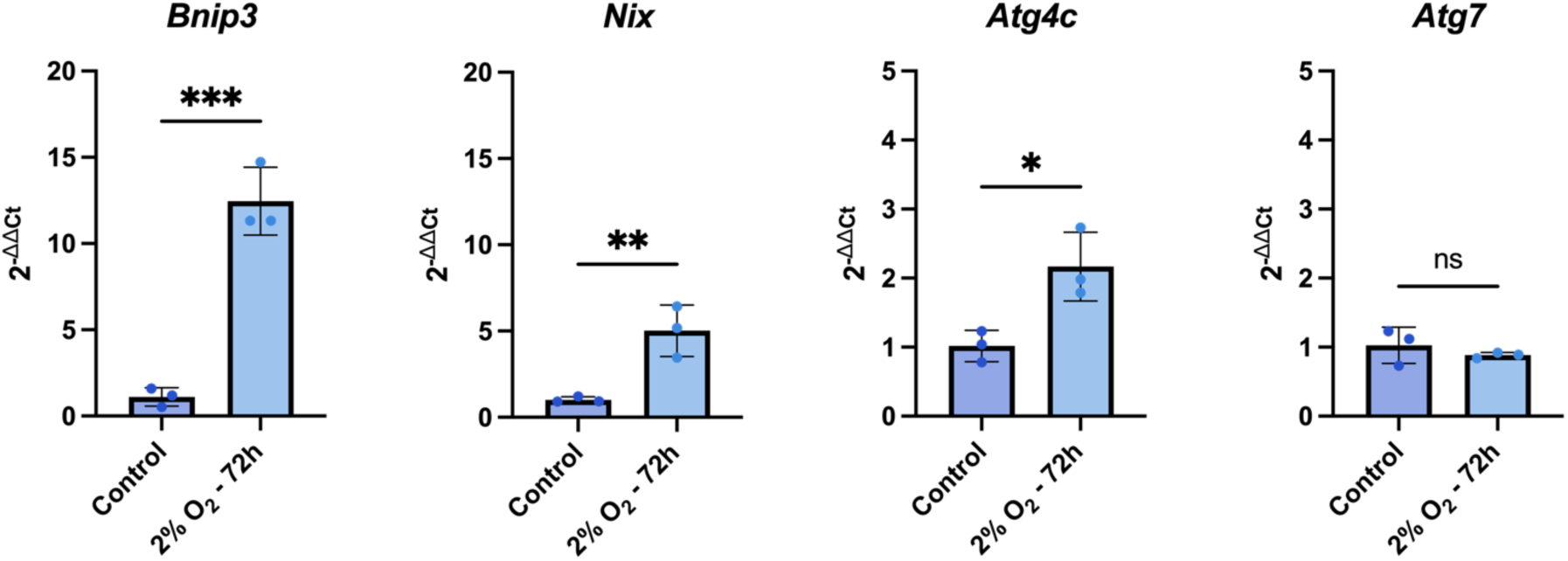
Hypoxic culture upregulates the mRNA levels of mitophagy-associated genes in CPCs. *Bnip3*, *Nix*, *Atg4c* and *Atg7* mRNA levels in CPCs cultured in hypoxia, in comparison to CPCs cultured in normoxia. Analysed using a t-test (n=3).

**Figure S3.**
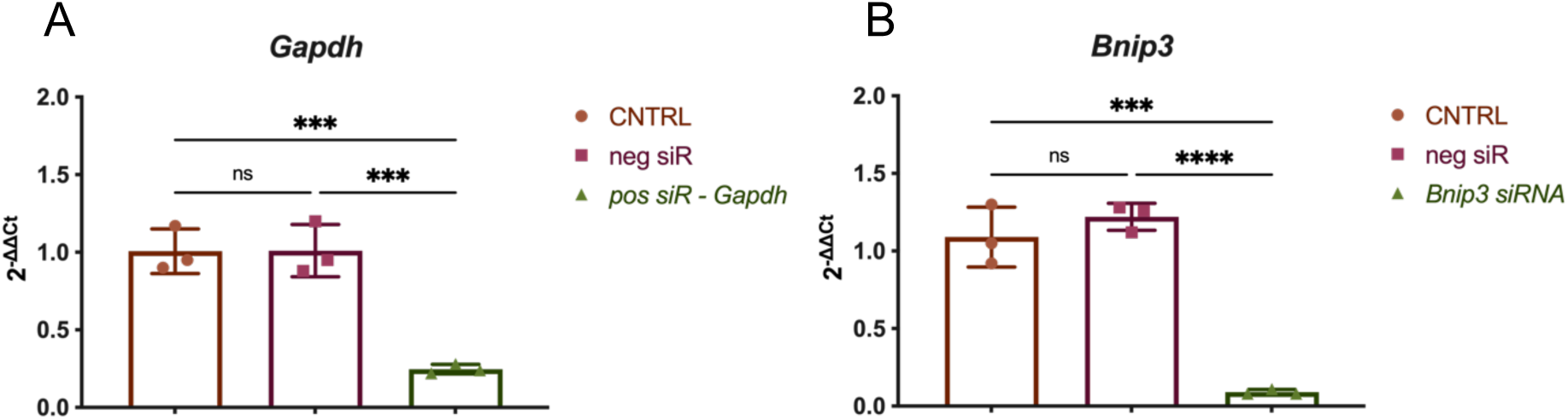
Validation of *Bnip3* siRNA. **(A)** *Gapdh* mRNA levels following transfection of CPCs with a positive control *Gapdh* siRNA. **(B)** *Bnip3* mRNA levels following transfection of CPCs with *Bnip3* siRNA at 100nM in comparison to a negative siRNA and an untransfected control. Analysed using a one-way ANOVA with Tukey’s multiple comparisons test (n=3).

**Figure S4.**
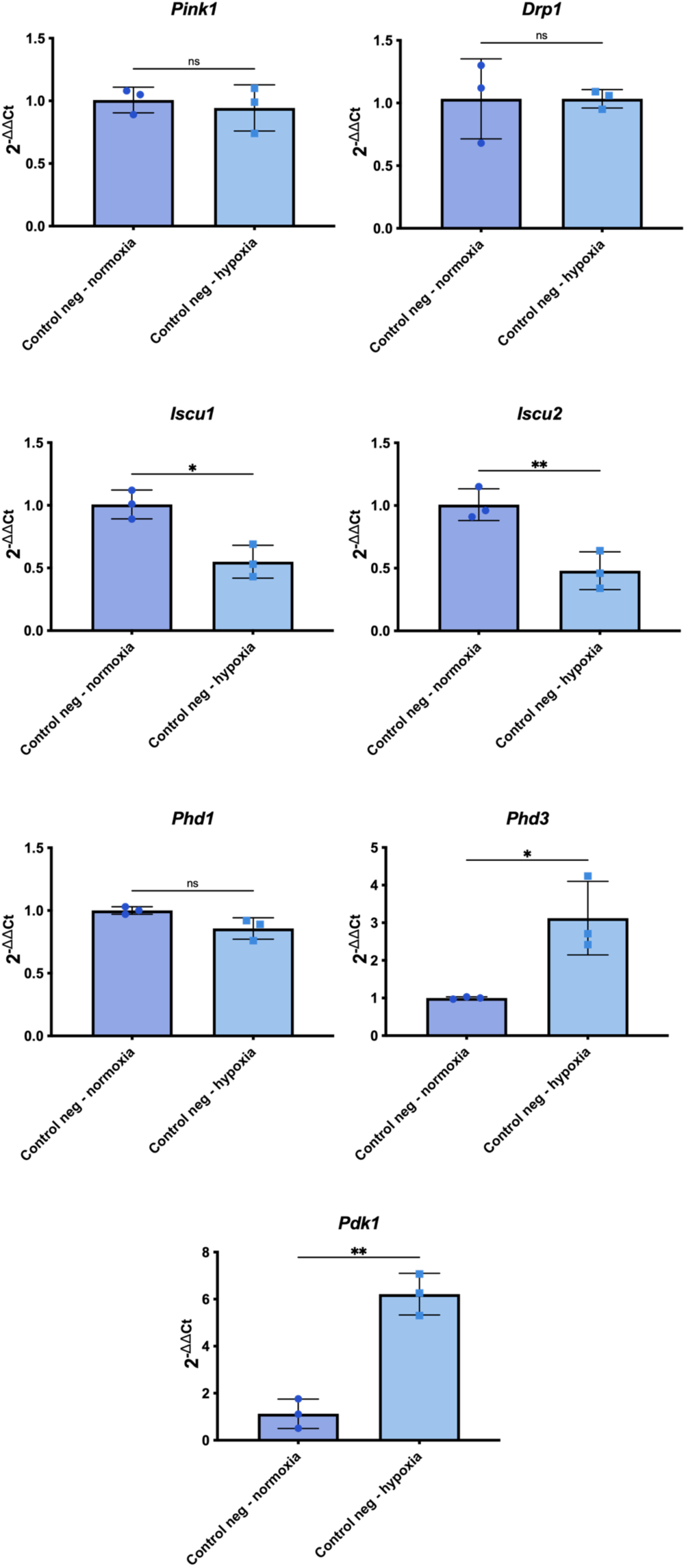
Expression of mitophagy- and hypoxia-associated genes in CPCs cultured in hypoxia. mRNA levels of mitophagy- and hypoxia-associated genes in CPCs transfected with a negative miRNA and cultured in hypoxia, in comparison to CPCs transfected with a negative miRNA and cultured in normoxia. The data are recycled from previous figures and were therefore added to the appendix. Analysed using a t-test to determine statistical significance (n=3).

**S.Table 1.**
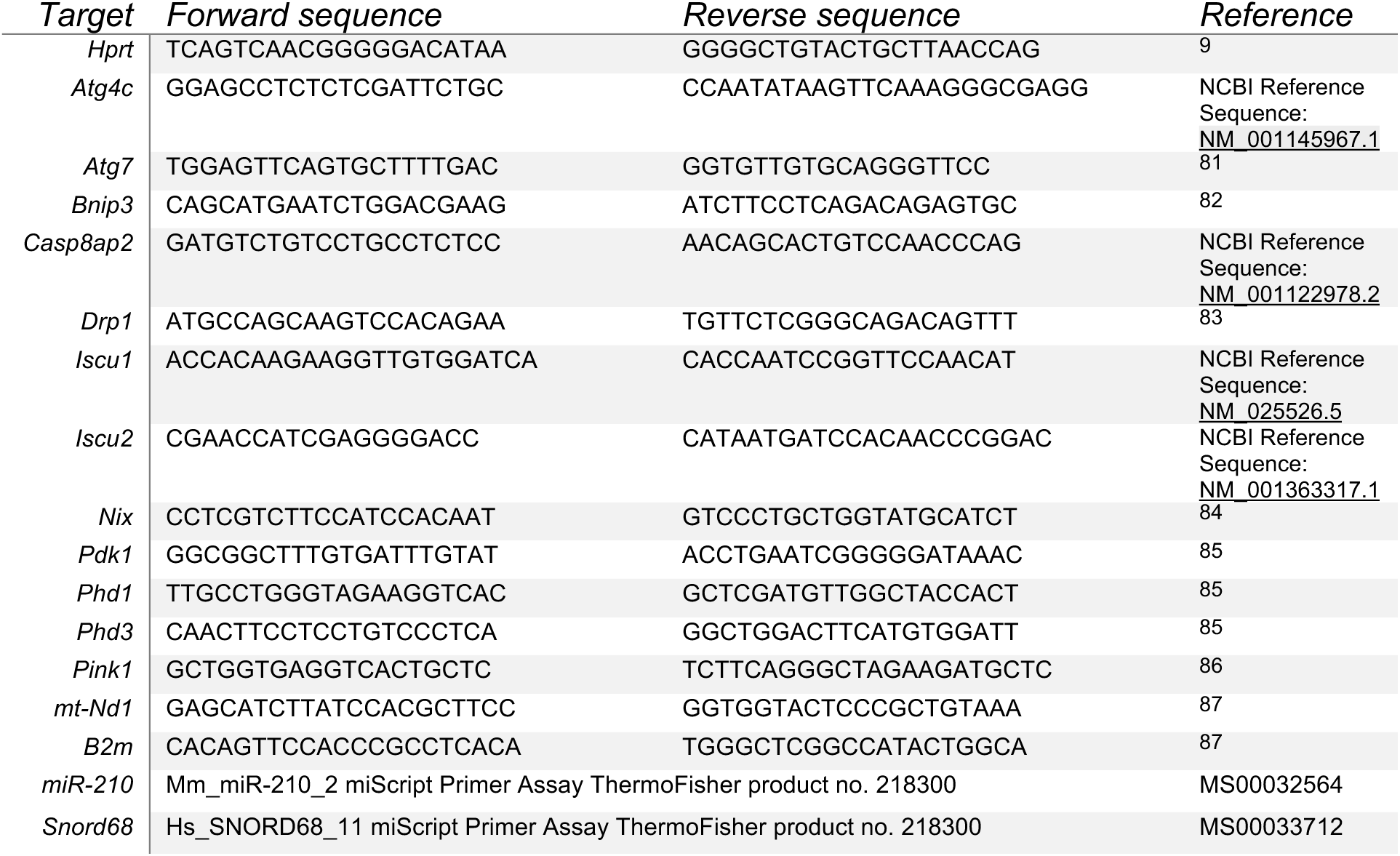
qRT-PCR primers.

